# A CRISPR activation screen identifies FBXO22 as an E3 ligase supporting targeted protein degradation

**DOI:** 10.1101/2023.09.15.557708

**Authors:** Ananya A. Basu, Chenlu Zhang, Isabella A. Riha, Assa Magassa, Felicia Ko, Xiaoyu Zhang

## Abstract

Targeted protein degradation (TPD) represents a potent chemical biology paradigm that leverages the cellular degradation machinery to pharmacologically eliminate specific proteins of interest. Although multiple E3 ligases have been discovered to facilitate TPD, there exists a compelling requirement to diversify the pool of E3 ligases available for such applications. This expansion will broaden the scope of potential protein targets, accommodating those with varying subcellular localizations and expression patterns. In this study, we describe a CRISPR-based transcriptional activation screen focused on human E3 ligases, with the goal of identifying E3 ligases that can facilitate heterobifunctional compound-mediated target degradation. This approach allows us to address the limitations associated with investigating candidate degrader molecules in specific cell lines that either lack or have low levels of the desired E3 ligases. Through this approach, we identified a candidate proteolysis-targeting chimera (PROTAC), 22-SLF, that induces the degradation of FKBP12 when the FBXO22 gene transcription is activated. 22-SLF induced the degradation of endogenous FKBP12 in a FBXO22-dependent manner across multiple cancer cell lines. Subsequent mechanistic investigations revealed that 22-SLF interacts with C227 and/or C228 in FBXO22 to achieve the target degradation. Finally, we demonstrated the versatility of FBXO22-based PROTACs by effectively degrading another endogenous protein BRD4. This study uncovers FBXO22 as an E3 ligase capable of supporting ligand-induced protein degradation through electrophilic PROTACs. The platform we have developed can readily be applied to elucidate protein degradation pathways by identifying E3 ligases that facilitate either small molecule-induced or endogenous protein degradation.

## Introduction

Traditional small molecule drugs function by directly interfering with the activities of proteins. However, many proteins lack suitable functional sites for rational drug design, presenting challenges in targeting them with small molecules. Some proteins, such as oncogenic adaptor proteins (e.g., MYD88) or transcription factors (e.g., MYC), are even considered “undruggable”^1^. Another complexity arises from proteins containing multiple functional domains. In such cases, compounds exclusively binding to one domain might inadequately deactivate the protein (e.g., IRAK4^2^). A promising alternative approach involves the use of small molecules that guide proteins to the cellular machinery responsible for proteolytic degradation, leading to the complete removal of the protein^3^. This targeted protein degradation (TPD) strategy employs two types of small molecules: 1) heterobifunctional compounds, known as PROTACs (proteolysis-targeting chimeras), which link E3 ligase ligands to substrate ligands using a structurally variable linker^4^; and 2) monofunctional compounds that form a complex involving specific E3 ligases and neo-substrate proteins, referred to as molecular glues (e.g., immunomodulatory drugs or IMiDs^5,6^). Targeted protein degradation offers several advantages. It can transform inert protein-binding small molecules into functional protein degraders, therefore expanding the range of druggable proteins. Additionally, this approach can function catalytically^7^, potentially reducing the required drug concentrations for a therapeutic impact. A rapidly growing number of proteins have demonstrated susceptibility to ligand-induced degradation within cells^3^. Several anticancer drugs in the market or under clinical evaluation operate through this mechanism^4^. Despite these successful cases, only a limited subset of the 600+ human E3 ligases has been identified as supportive of targeted protein degradation^8,9^. Moreover, these E3 ligases exhibit unique or restricted substrate preferences^8,9^, which are not currently predictable or easily controllable. This underscores the imperative to uncover additional ligandable E3 ligases with distinct properties, which would unlock the full potential of targeted protein degradation as a valuable pharmacological strategy.

Various strategies have been employed to identify E3 ligases for the application of TPD. One approach involves the rational design of ligands based on the structural characteristics of E3 ligases and their recognition of degrons. For instance, the discovery of the VHL ligand^10^ exemplifies this strategy. These logically engineered ligands can then be adapted into heterobifunctional molecules to evaluate their potential in degrading specific target proteins. Another effective strategy involves chemical proteomics, aiming to pinpoint small molecules that selectively bind to E3 ligases. This approach has led to breakthroughs such as identifying CRBN as a target of thalidomide^11^, RNF114 as a target of nimbolide^12^, and DCAF1 as a target of MY-1B^13^. Subsequently, these ligands can be transformed into heterobifunctional molecules for TPD applications. An alternative successful approach, which we have previously reported, entails target-degradation screening of a specialized collection of heterobifunctional compounds. These compounds consist of a ligand that binds to the desired target coupled with broad-spectrum electrophilic fragments. This method resulted in the discovery of E3 ligases DCAF16 and DCAF11, which facilitate TPD by interacting with covalent PROTACs^14,15^. A central premise of this technique is to utilize fragment electrophiles to potentially engage endogenously expressed E3 ligases. Given the unique catalytic properties of TPD, even partial engagement of these ligases can catalyze target degradation, which can be measured through techniques such as Western blot analysis or quantitative proteomics. Nonetheless, it’s important to acknowledge a limitation of this approach: the unpredictable availability of E3 ligases in the tested cell lines. This implies that the pool of accessible E3 ligases for candidate PROTACs might be restricted. On one hand, distinct cell lines exhibit varying E3 gene expression profiles, suggesting potential missed opportunities for exploitable E3 ligases that remain unexpressed. On the other hand, the mere presence of an E3 ligase does not guarantee its active functionality. For example, low protein expression levels can hinder the E3 ligase’s effectiveness.

To overcome this limitation and comprehensively evaluate the candidate PROTAC’s potential to engage various human E3 ligases, we have developed a novel strategy based on CRISPR transcriptional activation screening. This method can be implemented within a single cell line and has the potential to assess the interaction capabilities of most, if not all, human E3 ligases with candidate PROTACs, thus facilitating targeted protein degradation. This strategy involves the development of a focused single guide RNA (sgRNA) library, which is tailored to target the promoter regions of entire human E3 ligase genes. By employing this library, we can effectively induce the expression of human E3 ligases within a specific cell line. This systematic approach provides a means to identify previously undiscovered E3 ligases suitable for targeted protein degradation. In this study, using this technique, we successfully identified an E3 ligase F-box protein 22 (FBXO22), which actively facilitated the degradation of FKBP12 by an electrophilic bifunctional compound designed to bind FKBP12. The application of the FBXO22-targeting ligand enabled us to create an additional PROTAC capable of degrading the protein BRD4. This innovative strategy not only expands the repertoire of E3 ligases available for targeted protein degradation but also demonstrates the utility of CRISPR transcriptional activation screens in efficiently assessing candidate PROTAC interactions with a diverse range of human E3 ligases.

## Results

### A focused CRISPR-Cas9 transcriptional activation screen identifies an E3 ligase FBXO22 that supports electrophilic PROTAC-induced degradation of FKBP12

We utilized FKBP12 as the substrate protein for our degradation study. FKBP12 is a prolyl isomerase that has been frequently employed to investigate ligand-induced protein degradation^16,17^. To establish CRISPR-Cas9 transcriptional activation cells for the exploration of FKBP12 degraders, we initially transduced HEK293T cells with lentivirus carrying FKBP12-EGFP and subsequently sorted GFP+ cells (**Fig. 1a** and **Supplementary Fig. 1a**). The incorporation of a fluorescent tag allowed us to employ fluorescent intensity as a measurable parameter to assess FKBP12 protein levels. We first confirmed the degradation of FKBP12-EGFP using Len-SLF (**Supplementary Fig. 1b,c**), a PROTAC targeting FKBP12 through recruiting CRBN^15^. In contrast, the FKBP12 binder alone, SLF (synthetic ligand of FKBP), did not induce degradation of FKBP12-EGFP (**Supplementary Fig. 1b,c**). This data indicates the feasibility of using FKBP12-EGFP fusion protein to investigate FKBP12 targeting degraders. Subsequently, in FKBP12-EGFP-expressing cells, we stably expressed both dCas9-VP64 and MS2-P65-HSF1 (**Fig. 1a** and **Supplementary Fig. 1d**). This second-generation CRISPR-Cas9 transcriptional activation system incorporates the transcriptional activator VP64 along with additional coactivators p65 and HSF1, yielding enhanced transcriptional activation capabilities across diverse genes^18^. To validate the efficacy of gene transcriptional activation, we selected a well-studied gene IL-1β and generated two constructs with single-guide RNA (sgRNA) sequences targeting IL-1β promoter regions^18^. Transduction of FKBP12-EGFP, dCas9-VP64, and MS2-P65-HSF1-expressing cells with both IL-1β sgRNAs resulted in a substantial elevation of IL-1β gene expression (**Supplementary Fig. 1e**), confirming the effectiveness of this cell system.

**Fig. 1.**
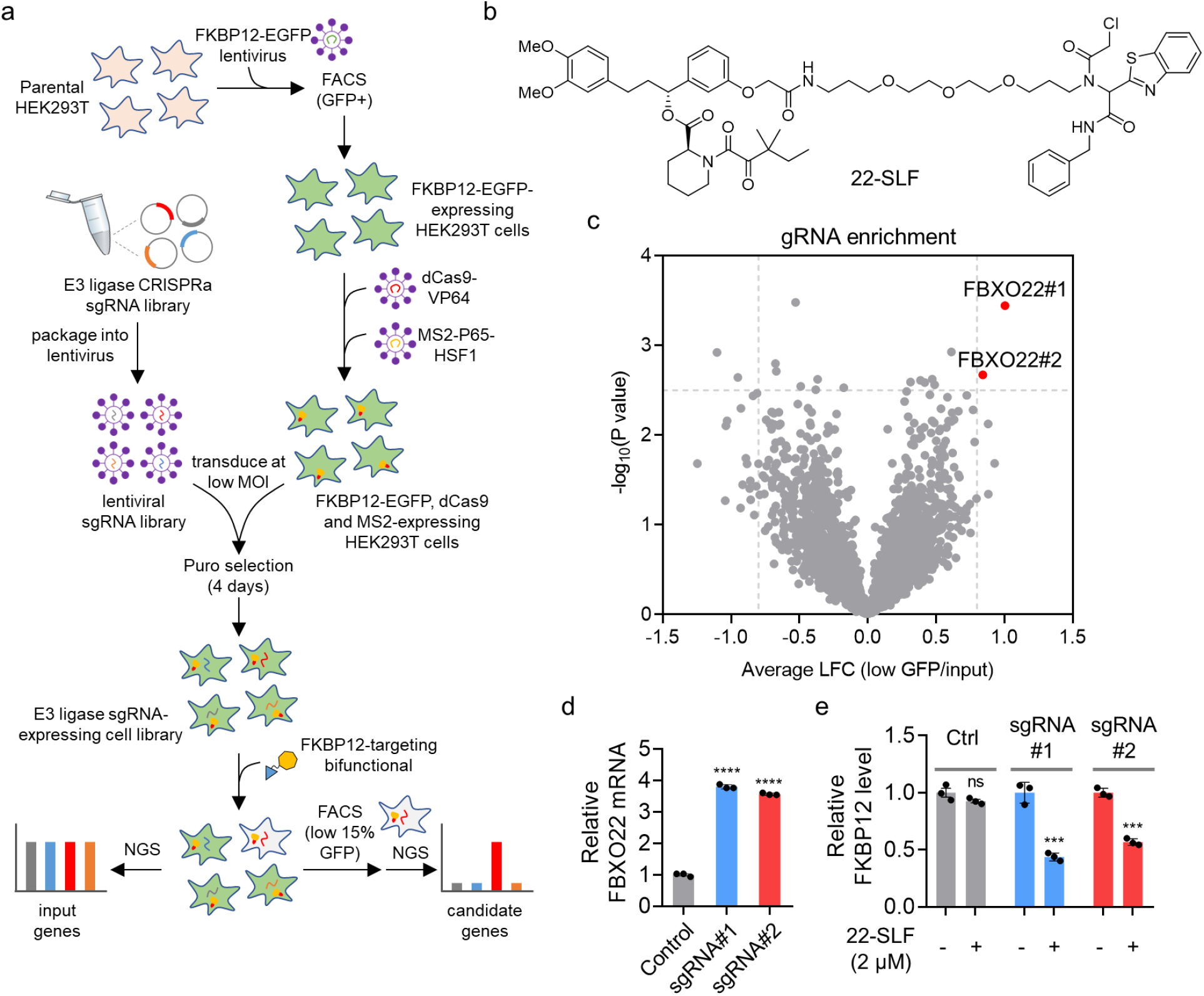
An E3 ligase focused CRISPR-Cas9 transcriptional activation screen identifies FBXO22 that supports 22-SLF-induced reduction in FKBP12-EGFP expression levels. **a**, Schematic representation of the steps in the CRISPR-Cas9 transcriptional activation screen. **b**, Structure of 22-SLF. **c**, Volcano plot showing the E3 ligase focused CRISPR-Cas9 transcriptional activation screen for FKBP12-EGFP degradation after treatment of 2 μM 22-SLF in HEK293T CRISPR-Cas9 transcriptional activation cells for 24 hours (n = 3 biological independent samples). **d**, Quantitative PCR analysis of FBXO22 mRNA levels subsequent to the transduction of sgRNAs targeting the promoter regions of the FBXO22 gene in HEK293T CRISPR-Cas9 transcriptional activation cells. The bar graph (n = 3 technical replicates) is representative of two independent experiments. The statistical significance was evaluated through unpaired two-tailed Student’s t-tests, comparing cells transduced with FBXO22 sgRNA to control sgRNA. Statistical significance denoted as \*\*\*\**P* < 0.0001 and ns: not significant. **e**, Fluorescence quantification of FKBP12-EGFP levels in HEK293T CRISPR-Cas9 transcriptional activation cells transduced with FBXO22-activating sgRNAs, subsequent to the treatment of 2 μM 22-SLF for 24 hours (n = 3 biological independent samples). The statistical significance was evaluated through unpaired two-tailed Student’s t-tests, comparing cells treated with 22-SLF to DMSO. Statistical significance denoted as \*\*\**P* < 0.001, \*\*\*\**P* < 0.0001 and ns: not significant.

With the CRISPR-Cas9 transcriptional activation cells, we aimed to conduct a CRISPR-Cas9 transcriptional activation pool screen to identify E3 ligases that support FKBP12 degradation by FKBP12-directed heterobifunctional compounds. To this end, we generated a focused sgRNA library containing 3,520 sgRNAs targeting 680 human E3 ligases (5 sgRNAs per E3 ligase, **Supplementary Table 1**) and packaged this library into lentivirus (**Fig. 1a**). The CRISPR-Cas9 transcriptional activation cells were transduced with the pooled sgRNA library at a multiplicity of infection (MOI) of approximately 0.3, yielding approximately 500 cells per sgRNA in each replicate. Following transduction, cells underwent puromycin selection (2 μg/mL) for a duration of 4 days. Subsequent to the selection process, we treated the cells with FKBP12-directed candidate PROTACs for 24 hours. The cells from the bottom 15% of the GFP population were isolated using fluorescence-activated cell sorting (FACS) (**Fig. 1a** and **Supplementary Fig. 2**). The relative abundance of sgRNAs in the sorted cells was then compared to the sgRNA distribution in the input cells prior to sorting.

Through an exploration of a series of FKBP12-directed heterobifunctional compounds, we identified a specific candidate, 22-SLF (**Fig. 1b)**, which exhibited substantial enrichment of two distinct sgRNAs targeting the same E3 ligase, FBXO22 (**Fig. 1c** and **Supplementary Table 2**). This discovery suggests that upon gene expression activation, 22-SLF transforms into a potent degrader, promoting the reduction in FKBP12-EGFP expression. To validate these findings, we individually introduced the two hit sgRNAs targeting FBXO22 promoter regions into the CRISPR-Cas9 transcriptional activation cells and confirmed their capability to activate FBXO22 mRNA expression (**Fig. 1d**). Subsequently, we subjected the FBXO22-activated cells to treatment with 22-SLF, which resulted in the loss of FKBP12-EGFP expression (**Fig. 1e**). This data was consistent with the findings from the CRISPR-Cas9 transcriptional activation screen.

### 22-SLF promotes FBXO22-dependent proteasomal degradation of FKBP12

Next, we sought to use Western blot analysis to measure the degradation of FKBP12 induced by 22-SLF. To this end, we stably overexpressed FLAG-tagged FKBP12 and HA-tagged FBXO22 in HEK293T cells, followed by subjecting the cells to varying concentrations of 22-SLF. 22-SLF-induced loss of FLAG-FKBP12 expression was FBXO22-dependent, as no reduction was observed in cells lacking HA-FBXO22 (**Fig. 2a**). Moreover, 22-SLF’s effect on FLAG-FKBP12 degradation was concentration-dependent (**Fig. 2a**). Notably, the loss of FLAG-FKBP12 expression can be blocked by the proteasome inhibitor MG132, as well as the neddylation inhibitor MLN 4924 and the FKBP12 ligand SLF (**Fig. 2a**). The intervention of MG132 in the rescue process suggests the proteasomal degradation of FLAG-FKBP12 induced by 22-SLF. The rescue enabled by MLN4924 indicates the involvement of a Cullin-RING ligase (CRL) in the degradation process^19^. This is consistent with the findings that FBXO22 belongs to SKP1-CUL1-F-box protein (SCF) ubiquitin ligase complexes, a subfamily of CRL^20^. In addition, the rescue by SLF demonstrates its competitive engagement with the FKBP12 binding site, indicating the indispensable role of 22-SLF’s binding to FKBP12 for the degradation process. Furthermore, the kinetics of 22-SLF-induced degradation of FLAG-FKBP12 are notably rapid, with nearly complete substrate degradation occurring within 2 hours (**Fig. 2b**). Collectively, this array of data supports the canonical mechanism of ligand-induced protein degradation. In this mechanism, 22-SLF orchestrates the recruitment of FBXO22 to FKBP12, resulting in the proteasomal degradation of FKBP12.

**Fig. 2.**
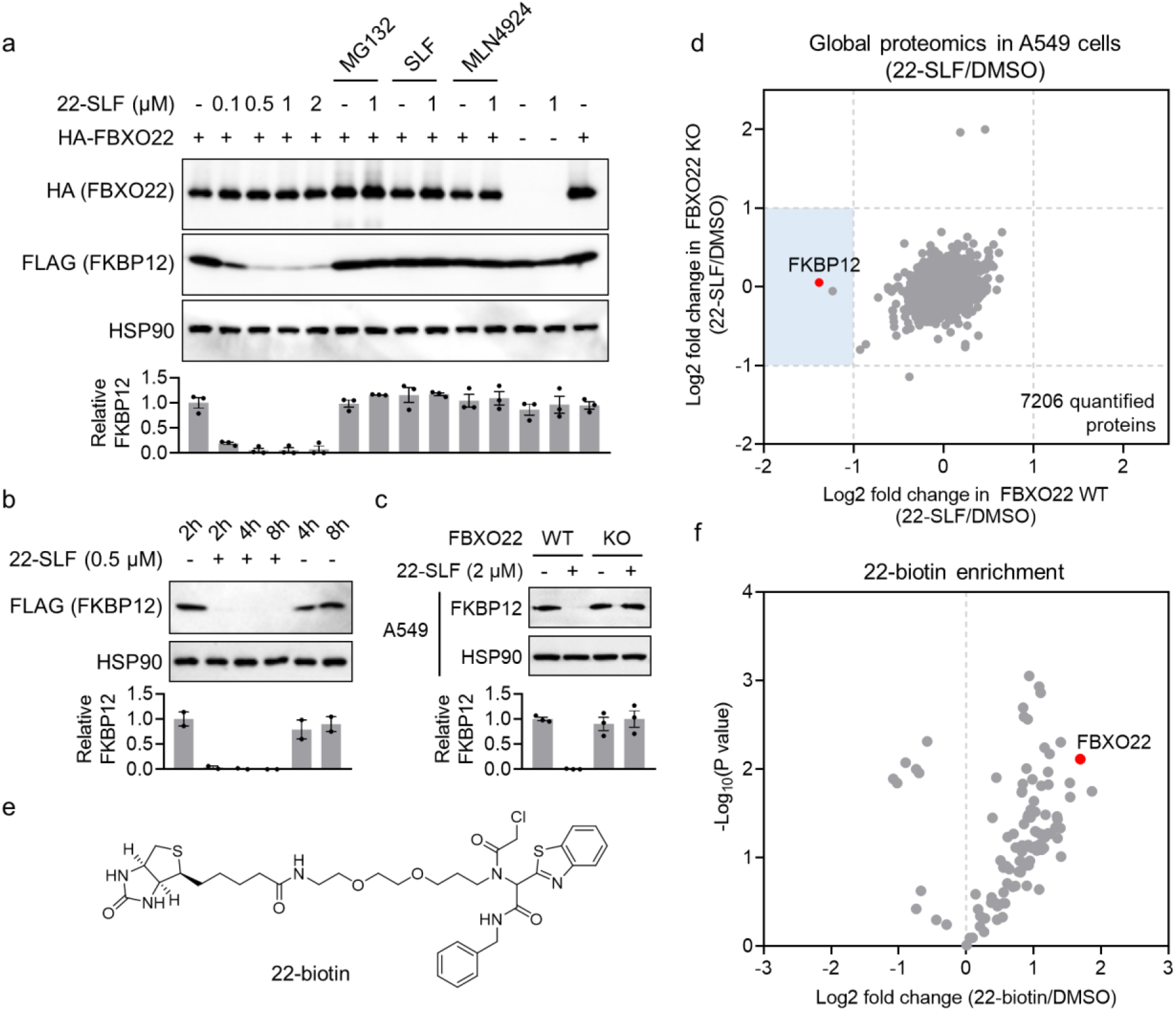
22-SLF promotes FBXO22-dependent proteasomal degradation of FKBP12. **a**, 22-SLF-induced FKBP12 degradation is dependent on FBXO22 and blocked by MG132 (1 μM), MLN4924 (1 μM) and SLF (25 μM) (n = 3 biological independent samples). The bar graph represents quantification of the FLAG-FKBP12/HSP90 protein content. Data are mean values ± SEM. **b**, Time-dependent degradation of FKBP12 by 22-SLF (n = 2 biological independent samples). The bar graph represents quantification of the FLAG-FKBP12/HSP90 protein content. Data are mean values ± SEM. **c**, 22-SLF (2 μM) induced FKBP12 degradation in A549 FBXO22 wildtype, but not knockout cells (n = 3 biological independent samples). The bar graph represents quantification of the FLAG-FKBP12/HSP90 protein content. Data are mean values ± SEM. **d**, Global proteomic analysis in A549 wildtype and FBXO22 knockout cells treated with 22-SLF (2 μM) for 24 hours (n = 2 biological independent samples for DMSO treatment, n = 3 biological independent samples for 22-SLF treatment). **e**, Structure of 22-biotin. **f**, 22-biotin pull-down with lysates of HA-FBXO22-expressing HEK293T cells followed by proteomic analysis revealed FBXO22 as one of the protein targets bound by 22-biotin (n = 2 biological independent samples).

As a F-box protein, FBXO22 has been demonstrated as a pivotal player in the context of tumor progression^21^. Its oncogenic functions come to the fore as it propels the ubiquitination and subsequent degradation of a diverse spectrum of substrates, such as histone lysine demethylase 4 subfamily KDM4A/B^22,23^, methylated p53^24^, p21^25^, PTEN^26^, KLF4^27^, and LKB1^28^. Further corroborating its relevance, FBXO22 demonstrates amplified expression within tumor tissues when juxtaposed with normal counterparts (**Supplementary Fig. 3a**). To establish the role of endogenous FBXO22 in facilitating 22-SLF-induced FKBP12 degradation, we employed FBXO22 knockout in a panel of cancer cell lines, including A549 (lung cancer), MDA-MB-231 (breast cancer), and PC3 (prostate cancer). The efficacy of FBXO22 knockout was verified through both genomic PCR and global proteomic analysis (**Supplementary Fig. 3b,c** and **Supplementary Table 3**). Notably, augmented expression of KDM4A and KDM4B – two substrates of FBXO22 – was evident in FBXO22 knockout cells, suggesting the functional disruption of FBXO22 in these cellular contexts (**Supplementary Fig. 3c**). Across all three cell lines (A549, MDA-MB-231 and PC3), 22-SLF promoted degradation of endogenous FKBP12, which was abolished upon FBXO22 knockout (**Fig. 2c** and **Supplementary Fig. 3d**). Furthermore, we conducted mass spectrometry (MS)-based global proteomic analysis on A549 parental and FBXO22 knockout cells, following the treatment with 2 μM 22-SLF for 24 hours. This experiment revealed the selective degradation of endogenous FKBP12 in FBXO22 wildtype, but not knockout context (**Fig. 2d, Supplementary Fig. 3d**, and **Supplementary Table 4**). Another protein, CUTA, was also observed to undergo reduction upon 22-SLF treatment in A549 parental cells, but not in the FBXO22 knockout setting (**Fig. 2d** and **Supplementary Fig. 3e**). To further substantiate the engagement of FBXO22 by the electrophilic portion of 22-SLF, we synthesized a biotin-conjugated probe 22-biotin (**Fig. 2e**). This probe was immobilized onto streptavidin-coated beads, which were subsequently subjected to interaction with lysates from HEK293T cells expressing HA-tagged FBXO22. Post-bead washing and on-bead trypsin digestion, the proteins associated with 22-SLF were analyzed using mass spectrometry, leading to the identification of FBXO22 as one of its binding targets (**Fig. 2f** and **Supplementary Table 5**).

### FBXO22 C227 and C228 are involved in 22-SLF-mediated degradation of FKBP12

The electrophilic α-chloroacetamide group in 22-SLF indicates a potential interaction with cysteine residues in FBXO22. To identify the specific cysteine residue(s) in FBXO22 engaged with 22-SLF, we treated HA-FBXO22-expressing HEK293T cells with either DMSO or 22-SLF (2 μM, 2 hours). Using competitive cysteine-directed activity-based protein profiling (ABPP), a chemical proteomic technology for the measurement of target engagement^29,30^, we sought to quantify the degree of blockade of iodoacetamide-desthiobiotin (IA-DTB)-modified cysteines on FBXO22. From a pool of 5,650 quantified IA-DTB-modified peptides, we identified four cysteines on FBXO22 (C83, C117, C228, and C365) (**Fig. 3a** and **Supplementary Table 6**), with approximately 20% of FBXO22 C228 being blocked by 22-SLF treatment (**Fig. 3b**), indicating the potential interaction with this cysteine by 22-SLF. Consequently, we mutated C228 to alanine and generated HA-FBXO22 C228A-expressing HEK293T cells (**Fig. 3c**). Interestingly, mutating C228 to alanine did not fully abolish 22-SLF-induced degradation of FKBP12, as the compound still partially degraded FKBP12 (**Fig. 3c**). Notably, in close proximity to C228 lies another cysteine, C227, implying the potential redundancy of these two cysteines in facilitating 22-SLF-induced FKBP12 degradation. Furthermore, the similarity in the fragmentation pattern of C227- and C228-modified peptides during MS analysis could imply that the engagement of C227 in FBXO22 by 22-SLF might not have been captured in the ABPP analysis. We then generated another single mutant (C227A) and a double mutant (C227AC228A) in FBXO22 and expressed these variants in HEK293T cells to assess 22-SLF-induced FKBP12 degradation. The data revealed that both single mutants, C227A and C228A, partially impeded 22-SLF-induced FKBP12 degradation, and the double mutant, C227AC228A, completely blocked the degradation of FKBP12 induced by 22-SLF (**Fig. 3c**). This suggests that C227 and C228 are likely to have overlapping roles in supporting 22-SLF-mediated FKBP12 degradation. To address the concern regarding potential conformational changes caused by mutations affecting the degradation, we conducted affinity purification-mass spectrometry (AP-MS) comparing the interactome of FBXO22 wildtype and C227AC228A double mutant. The results indicated that both forms were incorporated into the SKP1-CUL1-RBX1 E3 complex (**Supplementary Fig. 4a** and **Supplementary Table 7**), suggesting the likelihood of the proper folding of FBXO22 C227AC228A double mutant.

**Fig. 3.**
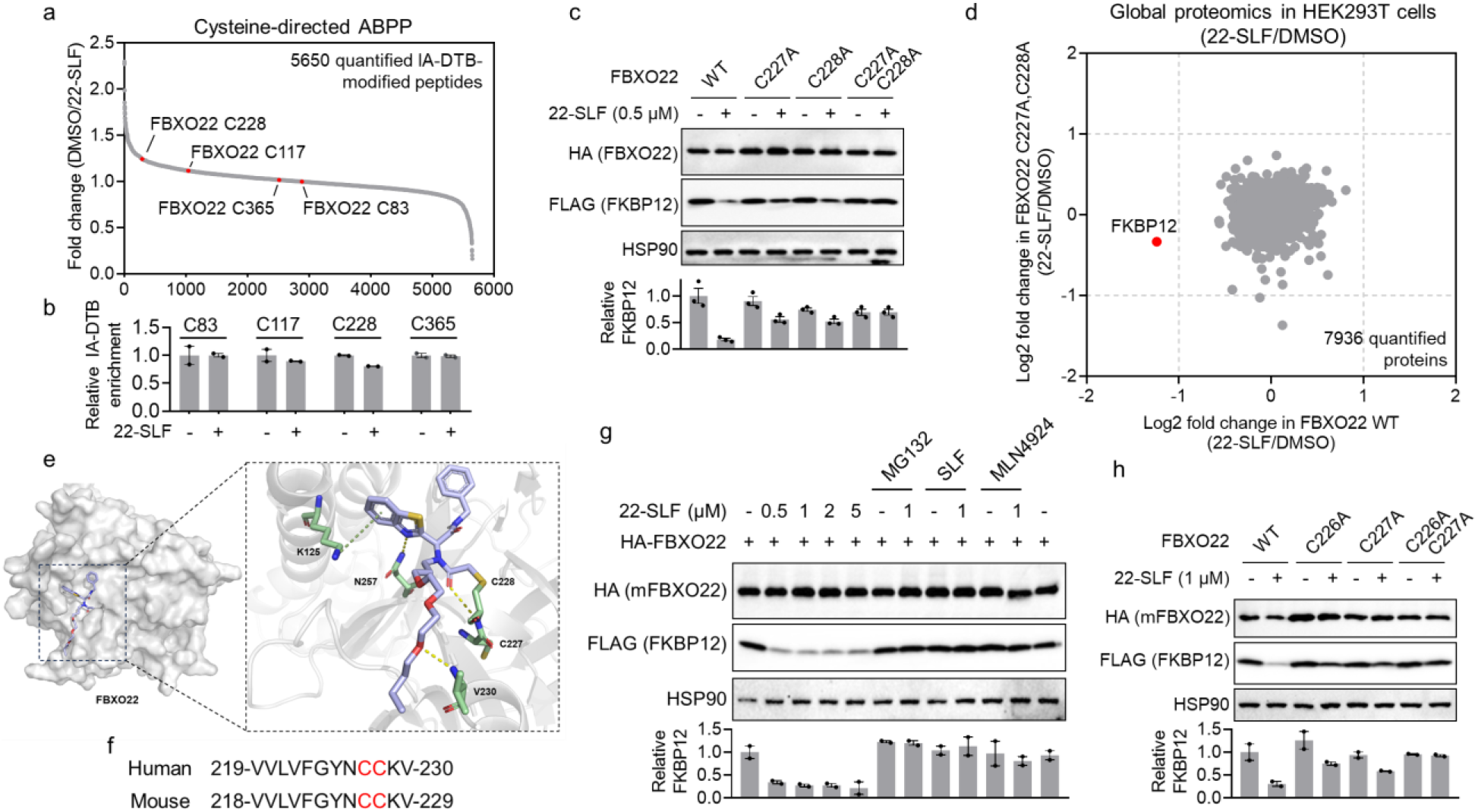
FBXO22 C227 and C228 are involved in 22-SLF-mediated degradation of FKBP12. **a**, Competitive cysteine-directed ABPP measured the degree of blockade of IA-DTB-modified cysteines by 22-SLF. Among 5,650 quantified IA-DTB-modified peptides, four cysteines were identified on FBXO22 (C83, C117, C228, and C365) (n = 2 biological independent samples). **b**, Quantification of four IA-DTB-modified peptides in FBXO22 treated with DMSO or 22-SLF (2 μM, 2 hours). **c**, Single mutation of C227 or C228 to alanine in FBXO22 partially blocked 22-SLF-induced degradation of FKBP12, while mutating both C227 and C228 to alanine in FBXO22 completely abolished the degradation (n = 3 biological independent samples). The bar graph represents quantification of the FLAG-FKBP12/HSP90 protein content. Data are mean values ± SEM. **d**, Global proteomic analysis in HEK293T cells expressing HA-FBXO22 wildtype or C227AC228A double mutant treated with 22-SLF (0.5 μM, 24 hours) (n = 2 biological independent samples for DMSO treatment, n = 3 biological independent samples for 22-SLF treatment). **e**, Modeling study reveals several hydrogen bond interactions between the electrophilic portion of 22-SLF and a pocket in FBXO22 involving C227 and C228. **f**, Sequence alignment of human FBXO22 (aa 219-230) and mouse FBXO22 (aa 218-229). **g**, Mouse FBXO22 supported 22-SLF-induced FKBP12 degradation. The degradation was blocked by MG132 (1 μM), MLN4924 (1 μM) and SLF (25 μM) (n = 2 biological independent samples). The bar graph represents quantification of the FLAG-FKBP12/HSP90 protein content. Data are mean values ± SEM. **h**, Mutation of both C226 and C227 in mouse FBXO22 abolished 22-SLF-induced degradation of FKBP12 (n = 2 biological independent samples). The bar graph represents quantification of the FLAG-FKBP12/HSP90 protein content. Data are mean values ± SEM.

Expanding on this, we performed global proteomic analysis in FBXO22 wildtype- and C227AC228A-expressing HEK293T cells, comparing the effects of 22-SLF treatment on the expression level of FKBP12. The findings revealed that among 7,936 quantified proteins, FKBP12 was the only protein degraded by 22-SLF in FBXO22 wildtype expressing cells, with no degradation observed in FBXO22 C227AC228A expressing cells (**Fig. 3d, Supplementary Fig. 4b** and **Supplementary Table 8**). Although the three-dimensional structure of FBXO22 remains undetermined, AlphaFold predictions demonstrate high confidence (pLDDT > 90) across most of the protein sequence, including the peptide encompassing liganded C227 and C228. To gain deeper insights into the binding model, we conducted a docking study focusing on the FBXO22 binding portion and the linker, with the predicted structure of FBXO22 generated by AlphaFold. The resulting binding model revealed a remarkable fit of this component on the surface of FBXO22 (**Fig. 3e**). Notably, the chloroacetamide warhead established a covalent bond with the C228 residue, consistent with findings from mass spectrometry analysis. Furthermore, hydrogen bonds were observed between the benzothiazole moiety and N257, the carbonyl group of the chloroacetamide and C227, as well as the oxygen atom of the PEG linker and V230. Additionally, the benzothiazole group engaged in a cation-pi interaction with K125, underscoring the potential of this scaffold as an effective ligand for the development of novel covalent PROTACs by engaging FBXO22.

FBXO22, a 403-amino acid protein, exhibits great conservation across mammals, with human and mouse FBXO22 sharing 93% identity (**Supplementary Fig. 5**). Intriguingly, the peptide sequence containing the two consecutive cysteines (C227 and C228) is identical between human and mouse (**Fig. 3f**). To explore whether mouse FBXO22 can facilitate 22-SLF-induced FKBP12 degradation and if the conserved cysteines (C226 and C227 in mouse FBXO22) are pivotal, we generated HEK293T cells expressing mouse FBXO22 wildtype, C226A, C227A, and C226AC227A. The results showed that 22-SLF treatment induced FBKP12 degradation in mouse FBXO22 wildtype-expressing cells, which can be blocked by MG132, SLF, and MLN4924 (**Fig. 3g,h**). Similar to human FBXO22, the single mutants, C226A and C227A, partially impeded degradation, while the double mutant, C226AC227A, fully blocked the degradation (**Fig. 3h**). This data underlines that mouse FBXO22 can integrate effectively into the human SKP1-CUL1-RBX1 complex and function as an E3 ligase supporting 22-SLF-induced FKBP12 degradation.

### 22-SLF induces a ternary complex between FKBP12 and FBXO22

It has been well established that the formation of ternary complex by degraders with their target protein and E3 ligase is the key step driving the subsequent target degradation^31^. To demonstrate the ternary complex involving 22-SLF, FKBP12 and FBXO22, we treated HEK293T cells expressing HA-FBXO22 and FLAG-FKBP12 with 22-SLF and a proteasomal inhibitor MG132. This experiment reveals that HA-FBXO22 co-immunoprecipitated with FLAG-FKBP12 in the presence of 22-SLF and MG132 (**Fig. 4a**), supporting a ternary complex formation involving 22-SLF, FBXO22, and FKBP12. Notably, the assembly of this ternary complex was partially hindered upon expression of FBXO22 C227A or C228A single mutant, and nearly completely abolished when the FBXO22 C227AC228A double mutant was introduced (**Fig. 4a**). This data further supports a model that 22-SLF engages C227/228 in FBXO22, leading to the formation of a ternary complex with and subsequent degradation of FKBP12.

**Fig. 4.**
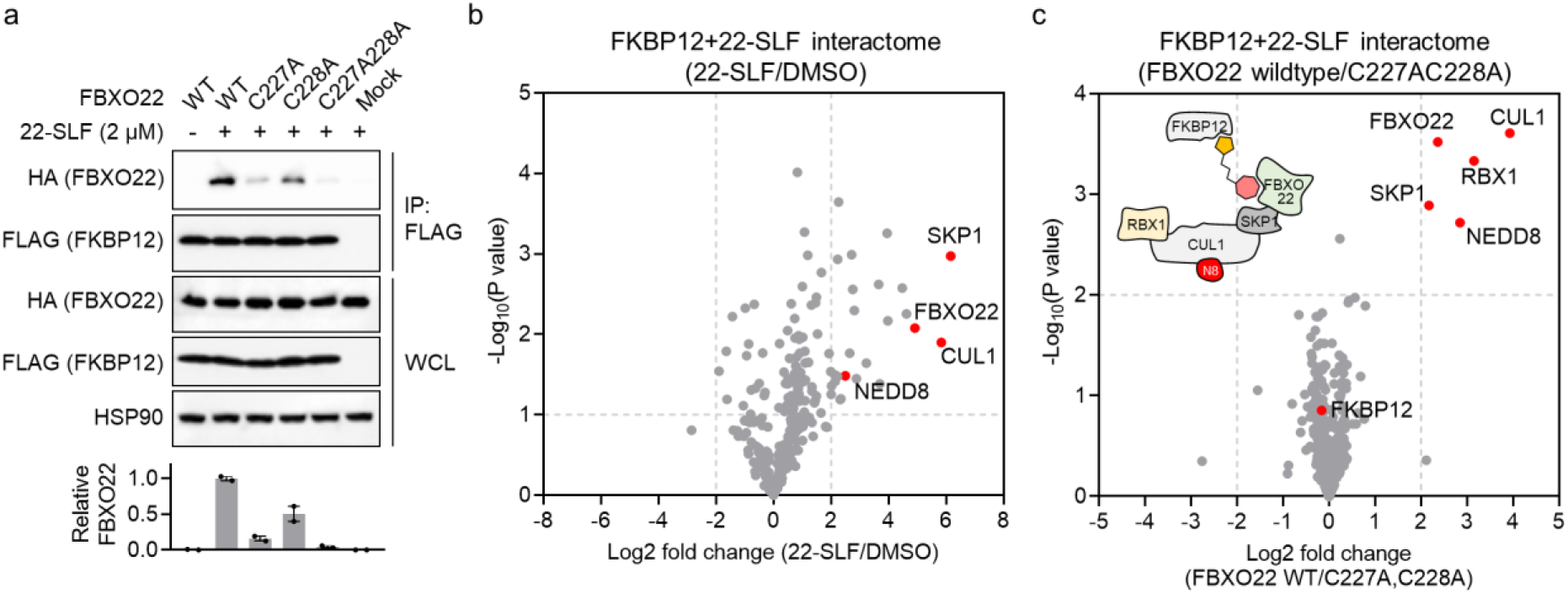
22-SLF induces the formation of a ternary complex involving 22-SLF, FKBP12 and FBXO22. **a**, A co-immunoprecipitation assay reveals that HA-FBXO22 wildtype, but not HA-FBXO22 C227AC228A double mutant, co-immunoprecipitated with FLAG-FKBP12 in the presence of 22-SLF and MG132 (n = 2 biological independent samples). The bar graph represents quantification of the immunoprecipitated HA-FBXO22 protein content compared to HA-FBXO22 protein in whole cell lysates (WCL). Data are mean values ± SEM. **b, c**, A FKBP12-directed enrichment proteomic analysis revealed that HA-FBXO22 wildtype and its associated components in the SKP1-CUL1-RBX1 E3 complex, but not HA-FBXO22 C227AC228A double mutant, co-immunoprecipitated with FLAG-FKBP12 in the presence of 22-SLF (2 μM) and MG132 (5 μM) (n = 2 biological independent samples).

To comprehensively assess the impact of 22-SLF on the FKBP12 interactome landscape, we employed an enrichment proteomic approach to identify proteins co-immunoprecipitating with FLAG-FKBP12 from HEK293T cells treated with 22-SLF. This analysis revealed several protein components of the FBXO22 complex, including FBXO22 itself, SKP1, CUL1, and NEDD8, as proteins recruited by 22-SLF (**Fig. 4b** and **Supplementary Table 9**). Furthermore, we conducted another enrichment proteomic study comparing FKBP12 interactome landscape in HEK293T cells expressing FBXO22 wildtype and C227AC228A double mutant. This experiment revealed the enrichment of the FBXO22 complex in FBXO22 wildtype-expressing cells, whereas the enrichment was absent in FBXO22 C227AC228A-expressing cells (**Fig. 4b** and **Supplementary Table 9**). Collectively, these findings indicate that 22-SLF effectively drives the formation of a ternary complex involving 22-SLF, FBXO22 and FKBP12 by engaging C227/228 in FBXO22, thereby facilitating the subsequent degradation of FKBP12.

### Harnessing FBXO22 for the degradation of additional protein targets

Finally, we sought to investigate the potential of FBXO22 to facilitate the degradation of a distinct protein. For this study, we opted to investigate BRD4 due to its significant role and the availability of a potent and selective ligand, JQ1, which has found extensive application in degrader development^16,32^. To explore this avenue, we synthesized a compound named 22-JQ1 by coupling the FBXO22 interacting electrophilic ligand to JQ1 (**Fig. 5a**). We performed a global proteomic analysis to assess the impact of 22-JQ1 on BRD4 degradation in A549 wildtype and FBXO22 knockout cells. Our findings revealed that 22-JQ1 induced a potent degradation of BRD4 in A549 wildtype, but not FBXO22 knockout cells (**Fig. 5c** and **Supplementary Table 10**). This result underscores the potential FBXO22 in the creation of additional PROTACs capable of degrading more protein targets.

**Fig. 5.**
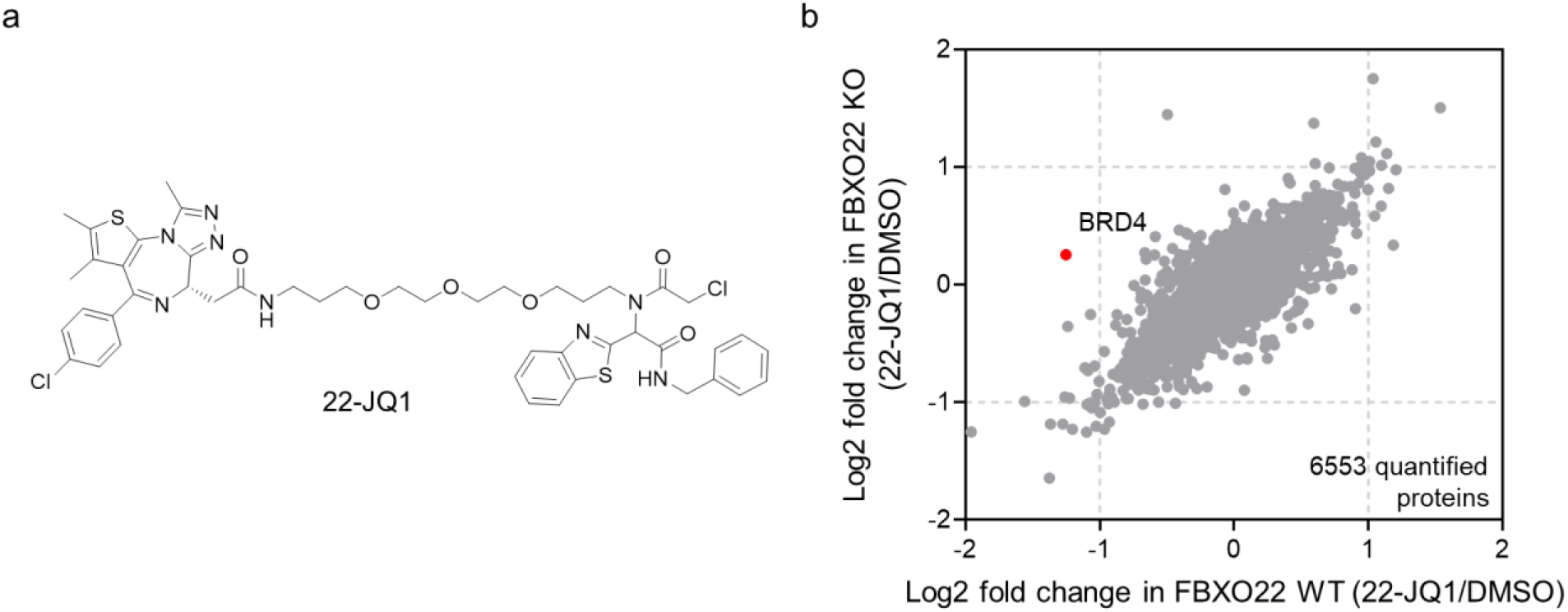
Harnessing FBXO22 for the degradation of BRD4. **a**, Structure of 22-JQ1. **b**, Global proteomic analysis in A549 wildtype and FBXO22 knockout cells treated with 2 μM of 22-JQ1 for 24 hours (n = 2 biological independent samples).

## Discussion

In this study, we employed a CRISPR-Cas9 transcriptional activation screen to unveil the capacity of the E3 ligase FBXO22 in supporting targeted protein degradation when engaged by electrophilic PROTACs. This novel approach holds promise for broad application, enabling the exploration of heterobifunctional compounds for their potential to degrade a diverse array of proteins of interest. Given the distinct gene expression profiles of various cell lines, candidate heterobifunctional compounds might exhibit a silent phenotype due to the absence of compatible E3 ligases. Our method offers an unbiased means of significantly boosting E3 ligase expression, facilitating confident assessment of candidate heterobifunctional compounds. This approach can be adaptable to both covalent and non-covalent compounds. Projecting forward, we are intrigued by the prospect of utilizing the screening approach reported here to uncover additional E3 ligases amenable to targeting by diverse PROTACs.

A commonly employed technique for identifying E3 ligases supportive of ligand-induced protein degradation is CRISPR-Cas9 knockout screens^33,34^. However, in instances where candidate compounds display moderate activity leading to partial target protein degradation, CRISPR-Cas9 knockout screens can yield high background noise, obscuring the deconvolution of specific E3 ligases. In such scenarios, our approach might offer an alternative solution. By activating E3 ligases, we can observe more pronounced target degradation, allowing for the enrichment of sgRNAs associated with the relevant E3 genes. Additionally, this technique holds value in cases where certain protein degraders exhibit degradation solely in specific cell lines or primary cells, which can pose challenges for establishing CRISPR-Cas9 knockout screens in hard-to-transduce cell models. In these situations, a CRISPR-Cas9 transcriptional activation screen could be established in transducible cells such as HEK293T cells. Absent activation of E3 ligase genes, the compound would exhibit inactive degradation behavior, but upon activating the target E3 ligase gene, the compound may become active, facilitating the isolation of cells where target degradation has occurred. Furthermore, this CRISPR-Cas9 transcriptional activation screen approach can serve to uncover novel endogenous degradation pathways, as well as identify redundant degradation pathways. By globally activating the expression of the entire E3 ligase family, we can evaluate the capabilities of all E3 ligases in degrading proteins of interest.

FBXO22 is a well-studied E3 ligase known for its involvement in tumor progression^21^. Given its elevated expression in tumors compared to normal tissue (**Supplementary Fig. 3a**), FBXO22 could offer an advantageous avenue for the degradation of cancer targets with potentially minimized toxicity in normal cells. Despite the extensive research into FBXO22’s biology, there remains a scarcity of studies focused on developing inhibitors or ligands for FBXO22. Our investigation establishes a foundational step in the development of FBXO22-targeting ligands with therapeutic potential. Our prior studies involving DCAF11 and DCAF16 have revealed that robust targeted protein degradation outcomes can be achieved with a low fractional engagement (< 40%) of the E3 ligases^14,15^. This attribute might provide a benefit to electrophilic PROTACs by enabling targeted protein degradation without significant perturbation of E3 ligase function. Similarly, our present study demonstrates that approximately 20% engagement of FBXO22 C227/C228 by 22-SLF effectively induces the degradation of FKBP12 (**Fig. 3b**). Collectively, these findings suggest that electrophilic PROTACs could achieve efficient degradation of target proteins with only partial engagement of E3 ligases, maintaining the physiological function of these ligases. Conversely, given the pivotal role of FBXO22 in tumor progression, inhibiting FBXO22 could hold promise as a cancer treatment strategy^21^. Thus, for cell contexts where FBXO22 inhibition could confer therapeutic benefits, potent ligands exhibiting high engagement with FBXO22 for degrader development could yield dual pharmacological effects – degradation of target proteins alongside FBXO22 inhibition.

It is important to note that our study currently presents an initial PROTAC leveraging FBXO22 to induce target degradation. We acknowledge that further optimization is required to enhance the potency and selectivity of electrophilic PROTACs acting through FBXO22, leading to the development of more advanced ligands. Additionally, since the electrophilic component of PROTACs contains a chiral center, future investigations will determine if these compounds display stereoselective degradation activity when evaluated as individual isomers. Exploring the potential for less reactive electrophiles (e.g., acrylamides) targeting FBXO22 would expand the breadth of ligand-induced protein degradation possibilities. Nonetheless, it’s promising that initial-generation electrophilic PROTACs acting through FBXO22 effectively promoted the degradation of multiple endogenous proteins. Our findings underscore the importance of integrating CRISPR activation screens, chemical proteomics, and covalent chemistry in PROTAC design, enabling the discovery of a broader spectrum of E3 ligases mediating ligand-induced protein degradation.

## Contributions

A.A.B. designed and conducted biochemical and cellular experiments to demonstrate target degradation; C.Z. conducted the modeling study; I.A.R. conducted experiments to demonstrate the effectiveness of GFP-based degradation platform; A.M., F.K., and X.Z. conducted the proteomics studies and performed data analysis; X.Z. supervised the project and, with contributions from all authors, wrote the manuscript.

## Acknowledgement

We gratefully acknowledge the support of the NIH (R00 CA248715, T32 GM105538, T32 GM149439), NSF GRFP, Damon Runyon Cancer Research Foundation (DFS-53-22), and the Illumina Pilot Project Program. We thank the Robert H. Lurie Comprehensive Cancer Center of Northwestern University for the use of the Flow Cytometry Core Facility.

## Materials and Methods

### Reagents

The anti-HA (clone#: C29F4, cat#: 3724), anti-FKBP12 (cat#: 55104), HRP-linked anti-HSP90 (clone#: C45G5, cat#: 79641), HRP-linked rabbit IgG (cat#: 7074) and HRP-linked mouse IgG (cat#: 7076) antibodies were purchased from Cell Signaling Technology. The anti-FLAG HRP antibody (clone M2, cat#: A8592), anti-FLAG affinity gel (clone M2, cat#: A2220), dCas9-VP64-blasticidin SAM CRISPRa helper construct 1 plasmid DNA (cat#: SAMVP64BST) and MS2-P65-HSF1-hygromycin SAM CRISPRa helper construct 2 plasmid DNA (cat#: SAMMS2HYG) were purchased from Sigma-Aldrich. Puromycin (cat#: ant-pr-1), blastcidin (cat#: ant-bl-05) and hygromycin (cat#: ant-hg-1) were purchased from InvivoGen. MG132 (cat#: S2619) was purchased from Selleck Chemicals. MLN4924 (cat#: 15217) and SLF (cat#: 10007974) were purchased from Cayman Chemical. Polyethylenimine (PEI, MW 40,000, cat#: 24765-1) was purchased from Polysciences, Inc. Enzyme-linked chemiluminescence (ECL) (cat#: 32106), ECL plus (cat#: 32132) western blotting detection reagents, Streptavidin agarose (cat#: 20349) and Tandem Mass Tag (TMT) isobaric label reagent (cat#: 90110) were purchased from Thermo Scientific. PureLink genomic DNA mini kit (cat#: K182001) was purchased from Invitrogen. FuGene 6 (cat#: E2692) transfection reagent and sequencing grade modified trypsin (cat#: V5111) were purchased from Promega. Cas9 endonuclease was purchased from Integrated DNA Technologies.

### Cell lines

HEK293T, A549, MDA-MB-231 and PC3 cells were obtained from ATCC. HEK293T and MDA-MB-231 cells were cultured in Dulbecco’s Modified Eagle Medium (DMEM, Corning) with 10% (v/v) fetal bovine serum (FBS, Omega Scientific) and L-glutamine (2mM, Gibco). A549 cells cultured in DMEM with 10% (v/v) FBS, L-glutamine (2mM) and MEM non-essential amino acids (Gibco). PC3 cells were cultured in F12K medium (Corning) with 10% (v/v) FBS. All the cell lines were tested negative for mycoplasma contamination.

### Generation of CRISPR-Cas9-mediated knockout cells

A549, MDA-MB-231 and PC3 cells with FBXO22 CRISPR-Cas9 knockout were generated through electroporation of Cas9-sgRNA ribonucleoprotein (RNP) complex using 4D-Nucleofector (Lonza Bioscience). Two sgRNAs targeting FBXO22 gene (FBXO22 sgRNA#1: GGACCCAUCGGAGCGUAACC; FBXO22 sgRNA#2: UCAACACGAAGGUGCUCCGC) were mixed for the electroporation. To confirm the knockout of FBXO22 gene, genomic DNAs from the electroporated cells were extracted using PureLink genomic DNA mini kit. FBXO22 gene was amplified by PCR and confirmed by DNA gel electrophoresis and DNA sequencing. Sequencing primers for FBXO22: TCCGAGCGTATTACGGAACG (forward) and AGACTGAACCACCAACCTGC (reverse).

### Cloning and Mutagenesis

Human and mouse FBXO22 cDNAs with N-terminal HA tag were purchased as gene block from Integrated DNA Technologies and cloned into pCDH-CMV-MCS-EF1-Blast vector (modified from pCDH-CMV-MCS-EF1-Puro vector by replacing PuroR with BlastR) via XbaI and EcoRI sites. FLAG-FKBP12 construct was generated as previously described^15^. FKBP12-EGFP construct was generated by cloning FKBP12 gene into a modified Artichoke plasmid (Addgene plasmid#: 73320) in which the selection marker puromycin was replaced with neomycin. FBXO22 mutants were generated using Q5 site-directed mutagenesis kit (New England Biolabs) following the manufacturer’s instruction.

### Library preparation of sgRNAs targeting human E3 ligases

sgRNAs targeting 680 human E3 ligases were designed using CRISPick (Broad Institute). 5 sgRNAs per E3 ligase were selected. sgRNAs with BsmBI restriction sites were ordered as oligonucleotide pool from Twist Bioscience. sgRNA library was amplified by PCR and purified using QIAquick gel extraction kit (Qiagen). CRISPRa vector was modified from LentiGuide-Puro (Addgene plasmid#: 52963) by inserting a tetraloop and stem loop sequences after the BsmBI restriction site^18^. Pooled DNA was cloned into CRISPRa vector in the presence of Esp3I (Thermo Scientific) and T7 ligase (New England Biolabs). The construct library was transformed into ElectroMAX Stbl4 competent cells (Invitrogen) using gene pulser Xcell electroporation system (Bio-Rad). The electroporated competent cells were cultured on Nunc Bioassay dish (Thermo Scientific). The construct library was extracted using plasmid Maxi kit (Qiagen).

### Generation of FKBP12 and FBXO22 stably expressed cells by lentivirus transduction

Lentivirus containing FLAG-FKBP12 and HA-FBXO22 were generated by co-transfection of FLAG-FKBP12/HA-FBXO22, psPAX2 and pMD2.G into HEK293T cells using FuGene 6 transfection reagent. Medium containing lentiviral particles were collected 48 hours post transfection, filtered with 0.45 μM Millex-HV sterile syringe filter unit (MilliporeSigma), and used to transduce HEK293T cells in the presence of 10 μg/mL polybrene. 48 hours post transduction, puromycin (2 μg/mL) or blastcidin (10 μg/mL) was added and incubated with the cells for 7 days.

### CRISPR-Cas9 transcriptional activation screen

Lentivirus containing sgRNA library targeting human E3 ligases was added to FKBP12-GFP, dCas9-VP64, and MS2-P65-HSF1-expressing HEK293T cells at a multiplicity of infection (MOI) of 0.3. 24 hours after transduction, cells were treated with 2 μg/mL puromycin for 4 days. After puromycin selection, cells were treated with 2 μM 22-SLF for 24 hours. Cells were sorted from the bottom 15% of the GFP population in BD FACSAria III flow cytometer. Total DNA was extracted using NucleoSpin blood mini kit (MACHEREY-NAGEL). sgRNA was amplified by PCR using mixed P5 primer and P7 primer with i7 index sequence. sgRNAs were quantified using the Illumina MiSeq. Read counts for sgRNAs targeting each gene were used to calculate fold changes and *P* values.

### Cell lysis and Western blot

Cells were lysed using RIPA lysis buffer (Thermo Scientific) containing 25 mM Tris-HCl, pH 7.6, 150 mM NaCl, 1% NP-40, 1% sodium deoxycholate, and 0.1% SDS. Lysis buffer was supplemented with the cOmplete protease inhibitor cocktail (Roche) before use. The cell suspension was subjected to sonication in 5 cycles of 40% power for 4 pulses each. Following sonication, the resulting mixture was centrifuged at 16,000g for 10 minutes at 4°C to obtain the supernatant. The protein concentration in the supernatant was quantified using the BCA assay (Pierce). The protein lysate was mixed with Laemmli sample buffer and heated at 95°C for 5 minutes. The proteins were separated using 4-20% Novex Tris-Glycine mini gels (Invitrogen) and transferred onto a 0.2 μM polyvinylidene fluoride (PVDF) membrane (Bio-Rad). The PVDF membrane was blocked with a solution containing 5% non-fat milk in TBST buffer (0.1% Tween 20, 20 mM Tris-HCl at pH 7.6, and 150 mM NaCl) for 1 hour at room temperature. Primary antibodies were diluted in 5% non-fat milk in TBST buffer (at a dilution of 1:5000 for FLAG and HA, and 1:1000 for others) and applied to the membrane. Incubation times were 1 hour at room temperature for FLAG, HA and β-Actin, and overnight at 4°C for others. Following antibody incubation, the membrane was washed three times with TBST buffer and incubated with a secondary antibody (diluted 1:5000 in 5% non-fat milk in TBST) for 1 hour at room temperature. After another three washes with TBST buffer, the chemiluminescence signal on the membrane was developed using ECL western blotting detection reagent (Pierce), and the resulting signal was captured using ChemiDoc MP (Bio-Rad). Relative band intensities were quantified using ImageJ.

### Real-time PCR analysis

HEK293T CRISPR transcriptional activation cells after sgRNA transduction were collected and total RNAs were extracted using the RNeasy mini kit (Qiagen). cDNAs were obtained by reverse transcription using the iScript cDNA kits (Bio-Rad). For real-time PCR analysis, SYBR Green real-time PCR master mix (Applied Biosystems) was used. Gene expression was quantified using QuantStudio 3 Real-Time PCR System (Applied Biosystems). sgRNA sequences for IL-1β gene transcriptional activation are: IL-1β sgRNA#1 TGGCTTTCAAAAGCAGAAGT, and IL-1β sgRNA#2 AAAAACAGCGAGGGAGAAAC. Primers for real-time PCR analysis are:

Forward primer of IL-1β: ATGATGGCTTATTACAGTGGCAA

Reverse primer of IL-1β: GTCGGAGATTCGTAGCTGGA

Forward primer of FBXO22: CGGAGCACCTTCGTGTTGA

Reverse primer of FBXO22: CACACACTCCCTCCATAAGCG

Forward primer of GAPDH: CTGGGCTACACTGAGCACC

Reverse primer of GAPDH: AAGTGGTCGTTGAGGGCAATG

### Immunoprecipitations

Cells were suspended and lysed in NP-40 lysis buffer (25 mM Tris-HCl at pH 7.4, 150 mM NaCl, 10% glycerol, 1% Nonidet P-40) supplemented with cOmplete protease inhibitor cocktail. The cell suspension was incubated on ice for 10 minutes. Subsequently, the mixture was centrifuged at 16,000 g for 10 minutes at 4°C, and the resulting supernatant was extracted for use in immunoprecipitation. For immunoprecipitation, FLAG or HA affinity gel (25 μL slurry per sample) was added to the protein lysates and rotated at 4°C for 2 hours. The affinity gel was then washed four times with immunoprecipitation washing buffer consisting of 0.2% NP-40, 25 mM Tris-HCl at pH 7.4, and 150 mM NaCl. Subsequently, the affinity gel was mixed with Laemmli sample buffer and heated at 95°C for 10 minutes. The resulting supernatant, containing the eluted proteins, was collected and used for subsequent western blot analysis.

### Mass spectrometry-based whole proteome analysis

Cells were lysed in 100 μL of PBS through sonication (10 pulses at 40% intensity, 3 rounds). Protein concentration was measured by a DC assay (Bio-Rad). 100 μg of proteins in 100 μL of lysis buffer were denatured using 8 M urea. For reduction, 5 μL of 200 mM DTT stock solution in water was added, and the mixture was heated to 65°C for 15 minutes. Alkylation was achieved by adding 5 μL of 400 mM iodoacetamide stock solution in water and incubating in dark at 37°C for 30 minutes. Subsequently, proteins were precipitated by adding 600 μL of MeOH, 200 μL of CHCl_3_, and 500 μL of water. After precipitation, protein pellets were washed with 1 mL of MeOH. The resulting protein pellets were solubilized in 160 μL of EPPS buffer (200 mM). 2 μg of LysC enzyme was added to each sample, and the digestion was carried out at 37°C for 2 hours. This was followed by the addition of 5 μg of trypsin to each sample for another round of digestion, which was allowed to proceed at 37°C for 12 hours. For TMT (tandem mass tag) labeling, 12.5 μg of resulting peptides in 35 μL of EPPS buffer were used. To each sample, 9 μL of CH_3_CN was added, and TMT tags (3 μL per sample, with a concentration of 20 μg/μL in CH_3_CN) were added. The samples were incubated at room temperature for 1 hour. The TMT labeling reaction was halted by adding 6 μL of a 5% hydroxylamine solution and 2.5 μL of formic acid. Subsequently, the samples were pooled together and subjected to fractionation followed by the analysis using liquid chromatography tandem mass-spectrometry (LC-MS) following the previously reported method^29^. Subsequently, the peptides were separated into 12 distinct fractions utilizing the Thermo Vanquish UHPLC fractionator. These resultant peptide fractions were then subjected to liquid chromatography tandem mass spectrometry analysis using an Orbitrap Eclipse Tribrid Mass Spectrometer coupled with a Vanquish Neo UHPLC System. The peptides were introduced onto an EASY-Spray HPLC column (C18, 2 μm particle size, 75 μm inner diameter, 250 mm length) and eluted at a flow rate of 0.25 μL/min, following the gradient: 5% buffer B (80% acetonitrile with 0.1% formic acid) in buffer A (water with 0.1% formic acid) from 0 to 15 minutes, 5% to 45% buffer B from 15 to 155 minutes, and 45% to 100% buffer B from 155 to 180 minutes. The nano-LC electrospray ionization source was set to a voltage of 1.5 kV. The analysis commenced with an MS1 master scan (Orbitrap analysis, resolution 120,000, m/z range 375-1600, RF lens 30%, standard AGC target, auto maximum injection time). In the MS2 analysis, precursor ions were quadrupole-isolated (isolation window 0.7) and then subjected to HCD collision in the ion trap (standard AGC, collision energy 32%, maximum injection time 35 ms). Following each MS2 spectrum, synchronous precursor selection (SPS) enabled the selection of up to 10 MS2 fragment ions for MS3 analysis. These MS3 precursors were fragmented by HCD and analyzed using the Orbitrap (collision energy 55%, AGC 250%, maximum injection time 200 ms, resolution 50,000). The RAW data was analyzed in Proteome Discoverer 2.5.

### Chemical proteomic analysis of cysteine reactivity in 22-SLF-treated cells

Cells were lysed in PBS through sonication (10 pulses at 40% intensity, 3 rounds). The protein concentration was determined using a DC assay and adjusted to 1 mg/mL. Subsequently, 500 μL of lysates were subjected to labeling with 100 μM iodoacetamide-desthiobiotin (IA-DTB) at room temperature for 1 hour. Protein precipitation was achieved by adding 500 μL of methanol and 100 μL of chloroform, followed by a methanol wash (1 mL). The resulting protein pellets were denatured using 90 μL of 9 M urea and 10 mM DTT in 50 mM tetramethylammonium bicarbonate. Alkylation was carried out using 50 mM iodoacetamide at 37°C for 30 minutes. Next, 350 μL of 50 mM tetramethylammonium bicarbonate was added to each sample, followed by the addition of 2 μg of trypsin. Digestion was allowed to proceed at 37°C for 12 hours. Subsequently, 50 μL of streptavidin-agarose beads were added to each sample and the mixture was gently rotated at room temperature for 2 hours. The beads were subjected to washing three times with 1 mL of washing buffer consisting of 0.2% NP-40, 25 mM Tris-HCl at pH 7.4, and 150 mM NaCl, three times with 1 mL of PBS, and two washes with 1 mL of water. Peptides were eluted using 300 μL of 50% acetonitrile containing 0.1% formic acid. The eluted peptides were subsequently dried using a SpeedVac vacuum concentrator. The subsequent steps of TMT labeling and LC-MS analysis were carried out following the methodology described above.

### Affinity purification–mass spectrometry

Cells were lysed in NP-40 lysis buffer comprising 25 mM Tris-HCl at pH 7.4, 150 mM NaCl, 10% glycerol, 1% Nonidet P-40, and the cOmplete protease inhibitor cocktail. Subsequent to centrifugation at 16,000 g for 10 minutes at 4°C, the supernatant was collected for immunoprecipitation. FLAG affinity gel (25 μL slurry per sample) was mixed with the protein lysates and incubated for 2 hours at 4°C. The gel was then washed four times using an immunoprecipitation washing buffer containing 0.2% NP-40, 25 mM Tris-HCl at pH 7.4, and 150 mM NaCl, followed by two washes with PBS. Elution of FLAG-FKBP12 and its associated proteins was carried out by heating the affinity gel at 65°C for 10 minutes in the presence of 8 M urea dissolved in PBS. The eluted proteins were subsequently reduced using 12.5 mM DTT at 65°C for 15 minutes, followed by alkylation with 25 mM iodoacetamide at 37°C for 30 minutes. The protein solution was diluted with PBS to reach a urea concentration of 2 M, and then digested using 2 μg of trypsin at 37°C for 12 hours. Subsequently, 6 μL of TMT tags (20 μg/μL in dry CH_3_CN) were added. The TMT labeling was allowed to proceed at room temperature for 1 hour, following which the reaction was quenched by the addition of 6 μL of a 5% hydroxylamine solution and 2.5 μL of formic acid. Subsequently, the samples were pooled and subjected to desalting using Sep-Pak C18 cartridge (Waters). The eluted peptide solution was dried using a SpeedVac vacuum concentrator. Peptides were then subjected to LC-MS analysis following the methodology described above.

### Streptavidin enrichment

Cells were lysed using NP-40 lysis buffer, which contained 25 mM Tris-HCl at pH 7.4, 150 mM NaCl, 10% glycerol, and 1% Nonidet P-40, supplemented with cOmplete protease inhibitor cocktail. The cell suspension was maintained on ice for 10 minutes. Following this, the mixture underwent centrifugation at 16,000 g for 10 minutes at 4°C, leading to the extraction of the resulting supernatant for subsequent use in enrichment. 22-biotin was incubated with streptavidin-agarose beads at room temperature for 2 hours. The beads were subjected to washing three times with 1 mL of washing buffer consisting of 0.2% NP-40, 25 mM Tris-HCl at pH 7.4, and 150 mM NaCl. Cell lysates were added to the probe-conjugated streptavidin-agarose beads and incubated at 4°C for 2 hours. The streptavidin-agarose beads were subjected to four washes using washing buffer, followed by two additional washes using PBS. Elution of interacted proteins was achieved by heating the streptavidin-agarose beads at 65°C for 10 minutes in the presence of 8 M urea dissolved in PBS. After elution, the proteins were subjected to reduction using 12.5 mM DTT at 65°C for 15 minutes, and subsequently alkylated using 25 mM iodoacetamide at 37°C for 30 minutes. To adjust the urea concentration to 2 M, the protein solution was diluted with PBS. The digestion was carried out using 2 μg of trypsin at 37°C for 12 hours. Following digestion, 6 μL of TMT tags (20 μg/μL in dry CH_3_CN) were added to the samples. The TMT labeling process was allowed to proceed at room temperature for 1 hour. Subsequently, the reaction was halted by adding 6 μL of a 5% hydroxylamine solution and 2.5 μL of formic acid. The samples were then pooled together and subjected to desalting using Sep-Pak C18 cartridge. The eluted peptide solution was dried using a SpeedVac vacuum concentrator. The dried peptides were subsequently prepared for LC-MS analysis following the methodology described above.

### Modeling study

The predicted crystal structure of FBXO22 from AlphaFold Protein Structure Database was utilized for the covalent docking. The protein preparation workflow in Maestro 13.4 (Schrödinger, LLC, New York, NY, 2022) was used to preprocess, optimize H-bond assignment, minimize energy and delete waters. Compounds were prepared by the LigPrep module with the OPLS4 force field. CovDock module was used for the docking process. The reaction type of nucleophilic substitution was selected, and the virtual screening approach was applied, along with default parameters for other settings. The final pose with low-energy conformation and good hydrogen-bond and cation-pi interaction geometries was selected. The figure was generated by PyMOL software.

### Statistical analysis

Quantitative data were depicted using scatter plots, displaying the mean accompanied by the standard error of the mean (SEM) represented as error bars. Differences between two groups were assessed using an unpaired two-tailed Student’s t-test. Significance levels were denoted as follows: \**P* < 0.05, \*\**P* < 0.01, \*\*\**P* < 0.001, and \*\*\*\**P* < 0.0001. Statistical significance was defined for *P* values < 0.05.

### Small Molecule Synthesis

22-SLF, 22-JQ1 and 22-biotin were synthesized by WuXi AppTec.

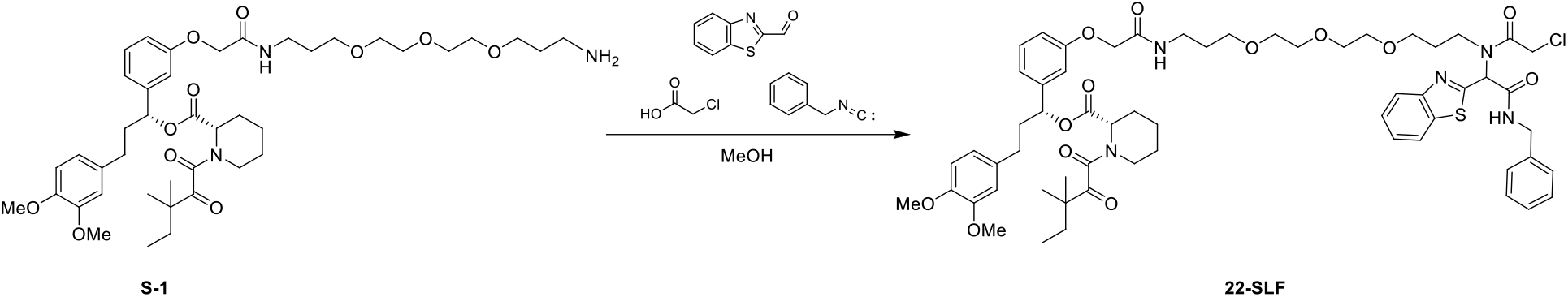

S-1 was prepared as reported previously. Benzothiazole-2-carboxaldehyde (21 mg, 0.13 mmol, 1.0 eq) was added to a solution of S-1 (105 mg, 0.13 mmol, 1.0 eq) in MeOH (0.3 mL, 0.5 M) and the reaction was stirred at room temperature for 12 h. After this, the benzyl isocyanide (19 mg, 0.16 mmol, 1.2 eq) and chloroacetic acid (18 mg, 0.16 mmol, 1.2 eq) were added and the reaction was stirred for 48 h. MeOH was removed and the residue was purified by preparative TLC to provide the desired analogue 22-SLF (35.8 mg, 0.031 mmol, 24%).

^**1**^**H NMR** (400 MHz, CDCl_3_) δ ppm 7.95-7.85 (m, 1H), 7.49-7.39 (m, 1H), 7.35-7.31 (m, 4H), 7.23-7.20 (m, 2H), 7.07-6.88 (m, 5H), 6.79-6.77 (m, 1H), 6.67 (s, 2H), 5.78 (s, 1H), 5.32 (d, J = 4.0 Hz, 1H), 4.72-4.62 (m, 1H), 4.48-4.45 (m, 3H), 4.39-4.14 (m, 3H), 3.86 (d, J = 2.8 Hz, 6H), 3.57-3.31 (m, 14H), 3.20-3.09 (m, 2H), 2.63-2.49 (m, 2H), 2.39-2.35 (m, 1H), 2.25-2.23 (m, 1H), 2.04-1.96 (m, 2H), 1.84-1.62 (m, 14H), 1.22 (d, J = 8.0 Hz, 6H), 0.89 (t, J = 8.0 Hz, 3H).

**HRMS** (ESI+) m/z calcd for C_60_H_77_ClN_5_O_13_S^+^ [M+H]^+^: 1142.4922, found 1142.4934.

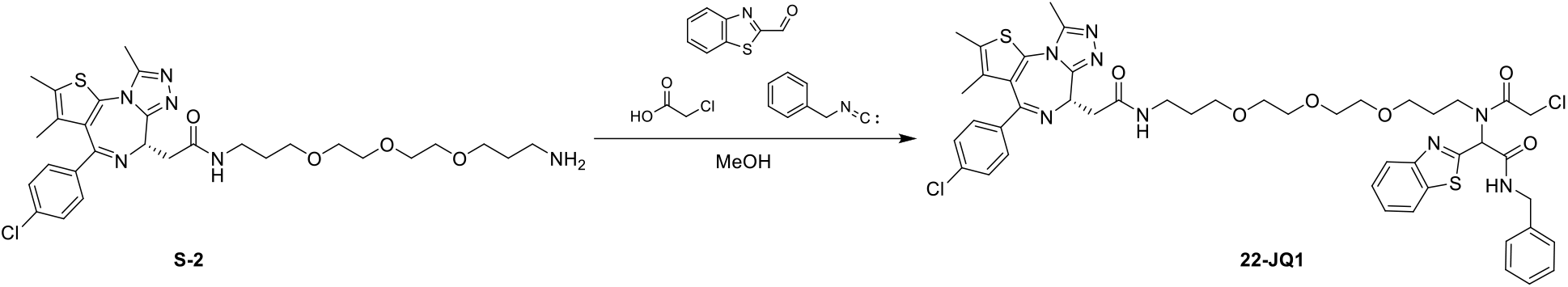

S-2 was prepared as reported previously. Benzothiazole-2-carboxaldehyde (65 mg, 0.40 mmol, 1.0 eq) was added to a solution of S-2 (240 mg, 0.40 mmol, 1.0 eq) in MeOH (0.8 mL, 0.5 M) and the reaction was stirred at room temperature for 12 h. After this, the benzyl isocyanide (56 mg, 0.48 mmol, 1.2 eq) and chloroacetic acid (54 mg, 0.48 mmol, 1.2 eq) were added and the reaction was stirred for 48 h. MeOH was removed and the residue was purified by preparative TLC to provide the desired analogue 22-JQ1 (48 mg, 0.05 mmol, 13%).

^**1**^**H NMR** (400 MHz, CDCl_3_) δ ppm 7.91-7.75 (m, 1H), 7.42-7.32 (m, 3H), 7.26-7.22 (m, 6H), 7.18-7.16 (m, 1H), 7.05-6.80 (m, 2H), 4.63-4.50 (m, 2H), 4.39-4.08 (m, 3H), 3.76-3.38 (m, 14H), 3.31-3.19 (m, 3H), 3.12-3.01 (m, 2H), 2.59-2.55 (m, 3H), 2.32 (s, 3H), 1.93-1.84 (m, 1H), 1.75-1.70 (m, 4H), 1.57(s, 3H), 1.24-1.18 (m, 1H).

**HRMS** (ESI+) m/z calcd for C_47_H_53_Cl_2_N_8_O_6_S_2_^+^ [M+H]^+^: 959.2901, found 959.2893.

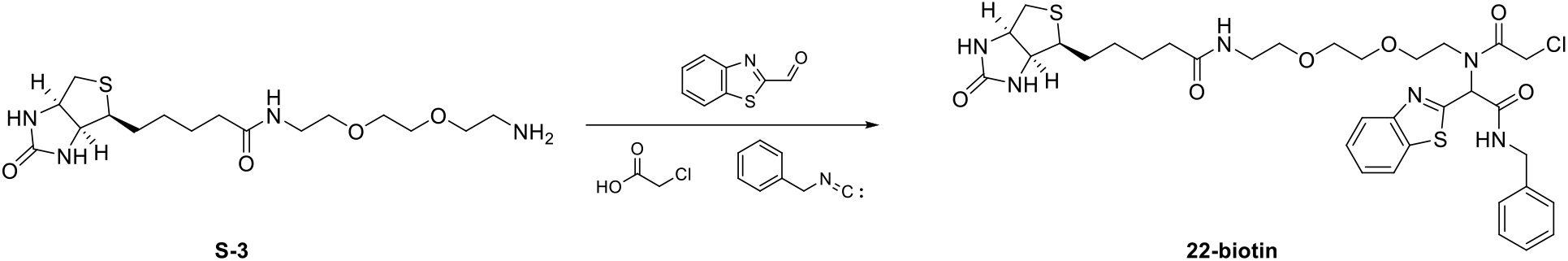

S-3 was prepared as reported previously. Benzothiazole-2-carboxaldehyde (175 mg, 1.07 mmol, 1.0 eq) was added to a solution of S-3 (400 mg, 1.07 mmol, 1.0 eq) in MeOH (2.0 mL, 0.5 M) and the reaction was stirred at room temperature for 12 h. After this, the benzyl isocyanide (149 mg, 1.28 mmol, 1.2 eq) and chloroacetic acid (144 mg, 1.28 mmol, 1.2 eq) were added and the reaction was stirred for 2 days. MeOH was removed and the residue was purified by preparative TLC to provide the desired analogue 22-biotin (65 mg, 0.089 mmol, 8%).

^**1**^**H NMR** (400 MHz, d_6_-DMSO) δ ppm 11.07 (s, 1H), 8.15-7.44 (m, 4H), 7.35-7.19 (m, 6H), 7.10-6.97 (m, 1H), 6.40-6.35 (m, 2H), 4.44-4.28 (m, 3H), 4.15 (d, J = 4.0 Hz, 1H), 4.12-4.09 (m, 1H), 3.87-3.78 (m, 1H), 3.59-3.51 (m, 3H), 3.38-3.23 (m, 7H), 3.13-3.05 (m, 3H), 2.80 (dd, J = 12.0, 8.0 Hz, 1H), 2.57 (d, J = 12.0 Hz, 1H), 2.05-2.02 (m, 2H), 1.62-1.39 (m, 4H), 1.33-1.21 (m, 3H).

**HRMS** (ESI+) m/z calcd for C_34_H_44_ClN_6_O_6_S_2_^+^ [M+H]^+^: 731.2447, found 731.2456.

**Supplementary Fig. 1.**
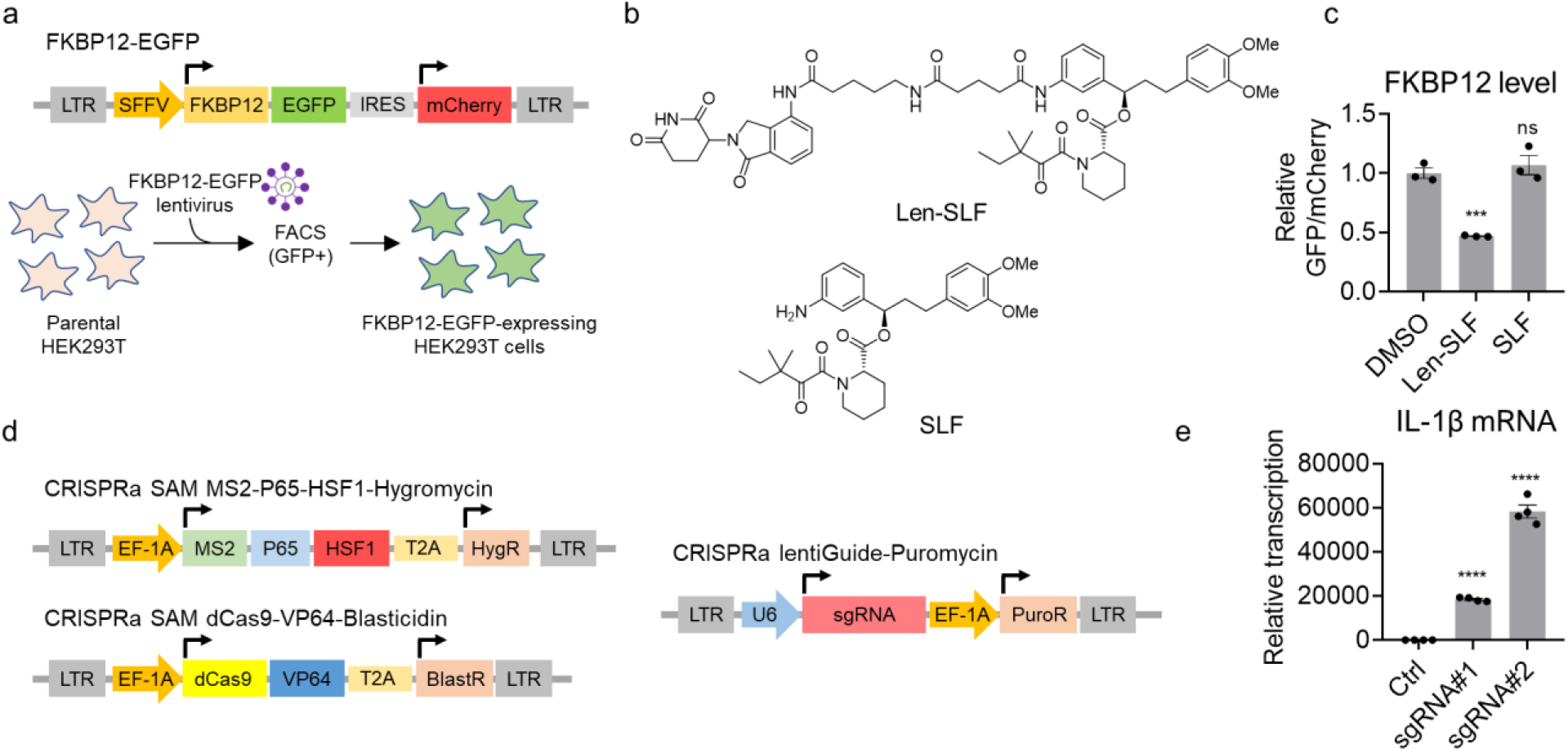
Generation of CRISPR-Cas9 transcriptional activation cells for the discovery of E3 ligases supporting ligand-induced degradation of FKBP12-EGFP. **a**, The construct of FKBP12-EGFP and a schematic representation of the generation of FKBP12-EGFP expressing HEK293T cells. **b**, Structures of Len-SLF and SLF. **c**, Fluorescence quantification of FKBP12-EGFP levels in HEK293T cells treated with 2 μM of Len-SLF or 20 μM of SLF for 24 hours (n = 3 biological independent samples). The statistical significance was evaluated through unpaired two-tailed Student’s t-tests, comparing cells treated with Len-SLF or SLF to DMSO. Statistical significance denoted as \*\*\**P* < 0.001 and ns: not significant. **d**, The constructs used for the CRISPR-Cas9 transcriptional activation screen. **e**, Quantitative PCR analysis of IL-1β mRNA levels subsequent to the transduction of sgRNAs targeting the promoter regions of the IL-1β gene in HEK293T CRISPR-Cas9 transcriptional activation cells. The bar graph (n = 4 technical replicates) is representative of two independent experiments. The statistical significance was evaluated through unpaired two-tailed Student’s t-tests, comparing cells transduced with IL-1β sgRNA to control sgRNA. Statistical significance denoted as \*\*\*\**P* < 0.0001.

**Supplementary Fig. 2.**
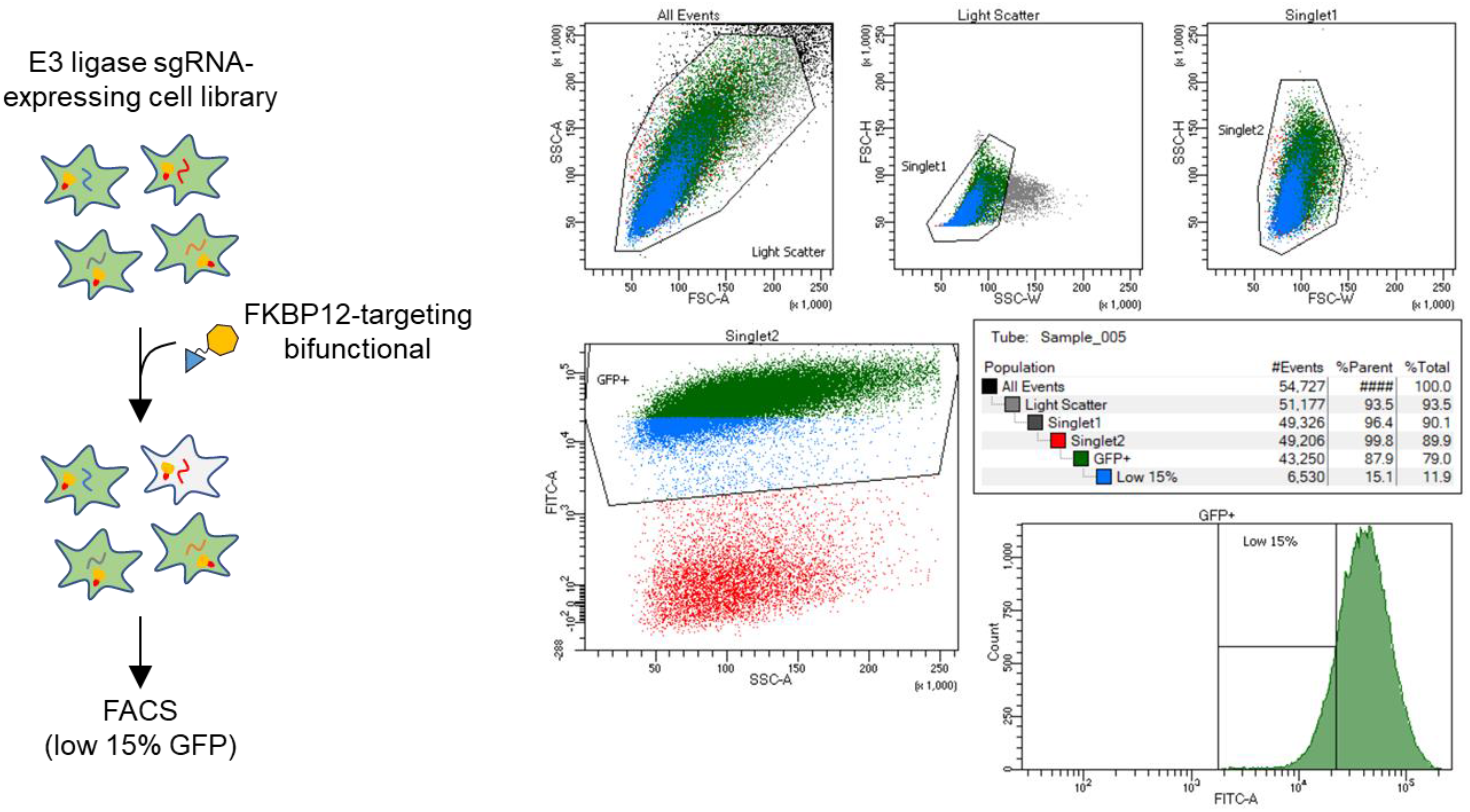
Procedure of fluorescence-activated cell sorting for the CRISPR-Cas9 transcriptional activation screen.

**Supplementary Fig. 3.**
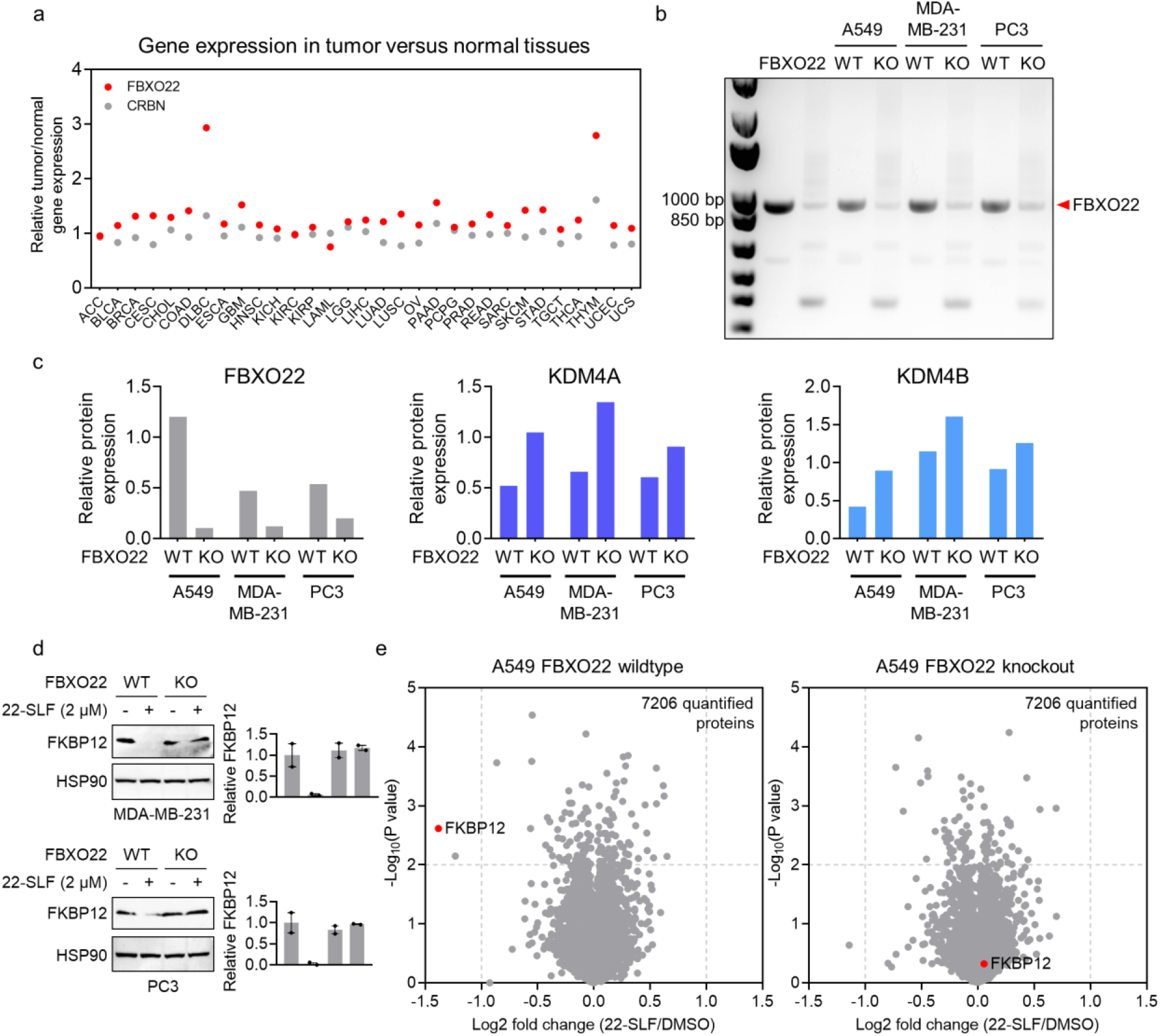
22-SLF promotes FBXO22-dependent proteasomal degradation of FKBP12. **a**, Gene expression ratio values of FBXO22 and CRBN between tumor and normal samples. Data is obtained from GEPIA (http://gepia.cancer-pku.cn/). **b**, Genomic PCR confirms FBXO22 knockout in A549, MDA-MB-231 and PC3 cells. **c**, Global proteomic analysis confirms FBXO22 knockout in A549, MDA-MB-231 and PC3 cells. KDM4A and KDM4B are two reported substrates of FBXO22. **d**, 22-SLF promoted reduction in FKBP12 levels in MDA-MB-231 and PC3 wildtype, but not FBXO22 KO cells (n = 2 biological independent samples). The bar graph represents quantification of the FLAG-FKBP12/HSP90 protein content. Data are mean values ± SEM. **e**, Global proteomic analysis in A549 wildtype and FBXO22 knockout cells treated with 22-SLF (2 μM, 24 hours).

**Supplementary Fig. 4.**
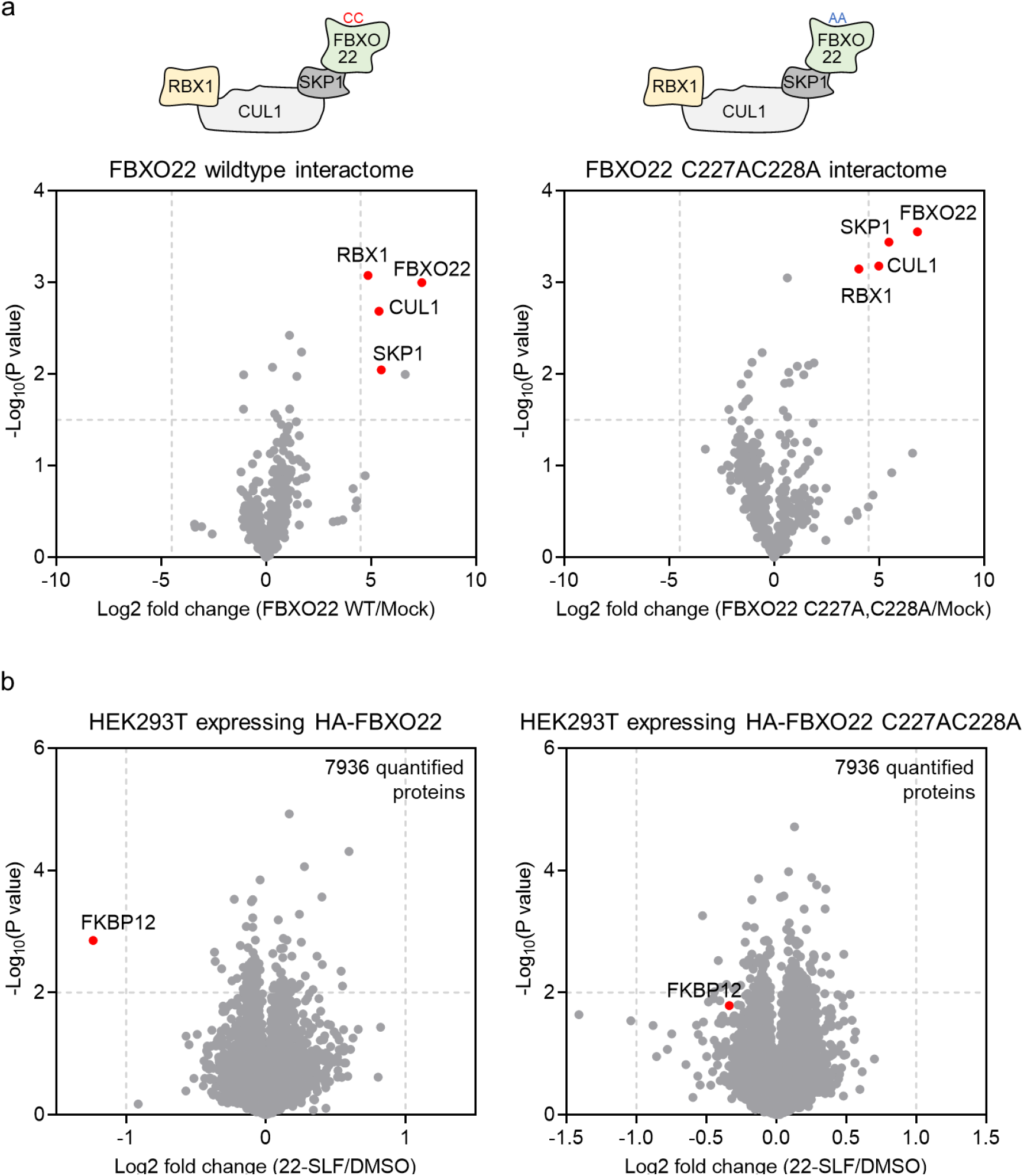
FBXO22 C227 and C228 are involved in 22-SLF-mediated degradation of FKBP12. **a**, Interactome studies of FBXO22 wildtype and C227AC228A double mutant revealed that both FBXO22 wildtype and C227AC228A double mutant were assembled into the SKP1-CUL1-RBX1 E3 complex. **b**, Global proteomic analysis in HEK293T cells expressing HA-FBXO22 wildtype or C227AC228A double mutant treated with 22-SLF (0.5 μM, 24 hours).

**Supplementary Fig. 5.**
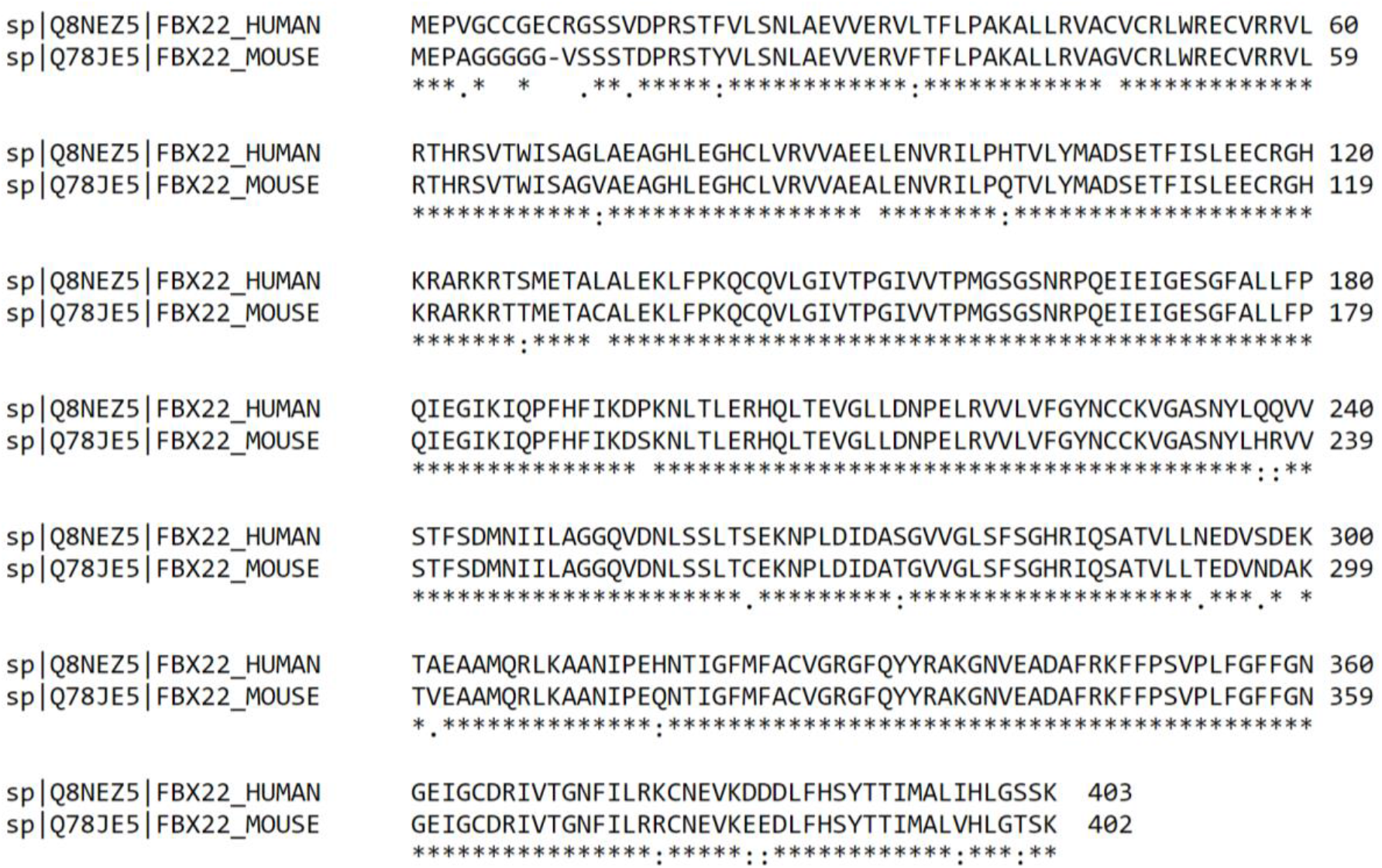
Sequence alignment of human and mouse FBXO22. Sequence alignment was performed in Clustal Omega.

